# Primary sensory neuron dysfunction underlying mechanical itch hypersensitivity in a Shank3 mouse model of autism

**DOI:** 10.1101/2024.12.29.630575

**Authors:** Damien Huzard, Mélanie Marias, Chloé Granat, Giulia Oliva, Vanessa Soubeyre, Gawain Grellier, Ahmed Negm, Jérôme Devaux, Emmanuel Bourinet, Amaury François

## Abstract

Autism Spectrum Disorder (ASD) is a neurodevelopmental disorder marked by social deficits, repetitive behaviors, and atypical sensory perception. The link between ASD and skin abnormalities, inducing itchiness, has never been investigated in depth. This study explores mechanical itch sensitivity in the Shank3^ΔC/ΔC^ mouse model. Key observations include heightened scratching in response to skin deformation and hypersensitivity to mechanical itch (i.e. alloknesis) in Shank3^ΔC/ΔC^ mice. In Shank3^ΔC/ΔC^ mice, *ex vivo* electrophysiological experiments revealed that C-fiber low-threshold mechanoreceptors (C-LTMRs) were hyporesponsive, and transcriptomic analysis showed a downregulation of TAFA4, a protein secreted by C-LMTRs. Interestingly, pharmacologically inhibiting Aβ-LTMR, important in mechanical itch initiation, abolished the itch hypersensitivity. Also, TAFA4 injections reduced the spontaneous scratching response to skin deformation but failed to restore itch sensitivity. Our data suggest that somatosensory deficits in Shank3^ΔC/ΔC^ mice lead to hypersensitivity to itchiness and indicate that two pathways might regulate mechanical itchiness, dependent on TAFA4.

## Introduction

Autism Spectrum Disorder (ASD) is a complex neurodevelopmental disorder characterized by persistent difficulties in social interactions, increased restricted and repetitive behaviors, as well as altered sensory perception [1]. In the absence of biological markers, ASD is currently diagnosed based on clinical scales focused on behavioral symptoms [1, 2]. Multiple clinical reports indicate that touch is often perceived differently in ASD patients [3, 4] and they commonly exhibit atypical sensory perception alterations [5, 6]. While some individuals with autism exhibit hyposensitivity to tactile stimuli [7], others display a marked intolerance to touch or hugging [8, 9] with sensory deficits differing across developmental stages. Despite these notable observations, the precise neuronal mechanisms underlying these sensory differences in autism are still not fully understood. In addition, it has been shown that there is a higher occurrence of skin-related disorders in ASD patients [10, 11], including atopic dermatitis and eczema [12, 13], as well as an increase in contagious itch responses [14, 15]. However, no clear relations between somatosensory defects and skin lesions in ASD have been identified or suggested yet.

Interestingly, various animal models employed to investigate and improve our understanding of ASD phenotypic differences have revealed peripheral and central somatosensory processing alterations. Recent studies demonstrated that the peripheral nervous system not only conveys tactile information but is also a key regulator of skin inflammatory responses and healing [16]. However, it remains to explore whether the peripheral nervous dysfunctions observed in ASD mouse models can be related to skin disorders [17–20]. In order to investigate if ASD mouse models can be utilized to study the mechanism behind ASD associated skin disorder, we focused on Shank3^ΔC/ΔC^ mice, which carry a specific gene mutation linked to Phelan–McDermid syndrome and ASD in humans [19]. The primary objective was to explore the effects of this mutation on the functioning of primary sensory neurons functions and tactile modalities, particularly focusing on abnormal mechanical itch in response to light punctate stimuli and skin deformation. This involved behavioral and pharmacological manipulations of somatosensory neurons, supplemented by *ex vivo* analyses of primary sensory neurons reactivity, and transcriptomic analysis of genes involved in itch and inflammatory responses.

In this study, we reveal that mice carrying the Shank3^ΔC/ΔC^ mutation display an increased scratching response to skin deformation and hypersensitivity to mechanical itch, accompanied by hypofunctional C low-threshold mechanoreceptors (C-LTMRs). Pharmacological studies revealed that the abnormal mechanical itch response depends on Aβ-LTMRs activity and mimicking C-LTMRs-induced analgesia, using an injection of TAFA4, reduces the spontaneous scratching behavior to skin deformation. We thus suggest that C-LTMRs are a key mediator of abnormal itch in Shank3^ΔC/Δ^ mice and propose a dual mechanism being dependent, or not, on TAFA4, in the control of the scratching behavioral response to mechanical skin challenges.

## Material and Methods

### Mice

All mice were kept on a C57BL/6J background by back-crossing with B6J mice for more than five generations. Heterozygous Shank3^ΔC/+^ male and female mice were bred to produce homozygous KO Shank3^ΔC/ΔC^ and WT Shank3^+/+^ littermates. Mice were housed 2-4 animals of similar genotype per cage in ventilated cages (Tecniplast, Italy). Cages were changed weekly, food and water was available *ad libitum*. All behavioral experiments were conducted in 8–20 weeks old animals. Shank3^ΔC^ mouse line was a gift from Dr. Julie Perroy.

### Ethics

All animal procedures complied with the welfare guidelines of the European Community and were approved by the local ethic committee, the Herault department Veterinary Direction, France and the French ministry for higher education, research and innovation (Agreement Numbers: 2017100915448101 and 2022120112076042).

### Sex of mice studied

Initial behavioral experiments were performed on animals from both sexes. Data from males and females were first compared and analyzed separately (Supplementary Fig. 2). No major difference was observed in behavioral outputs in the marble burying test (Supplementary Fig. 2a), in grooming (Supplementary Fig. 2b) digging (Supplementary Fig. 2c) or mechanical itch hypersensitivity (Supplementary Fig. 2d-f). Solely male mice were used for the qPCR, nerve physiological analysis and pharmacological manipulations, when a lower number of animals were tested.

### For all behavioral experiments

Prior to all behavioral experiments mice were moved in the experimental room at least 30 minutes before starting behavioral testing. Experiments were performed in the morning (from 8 am to 1 pm) under approximately 150-180 lux. The random assignment of mice to either the wild-type or knockout experimental groups was based on Mendelian inheritance. To blind the experimenters, each cage included in a cohort was attributed with one color at random. Then, in each cage, animals randomly received a number and on test days animals were placed in the test compartments so that all animals with the same number were placed side by side (e.g. one red, one blue, two red, two blue, etc…). When the experiment required multiple tests, the testing order was randomized daily, with each animal tested at a different time each test day. The experimenters who performed behavioral testing, recording, and analysis were blind to the animal genotype condition and/or treatment. The assignment of the color-coded labeling to the genotype and/or the pharmacological treatment was done by a co-experimenter after analysis. No mice were excluded from our experiments.

### Testing hind paw mechanical sensitivity with Von Frey filaments

Mice were first habituated for 45 min to the experimental boxes and grid for two consecutive days before testing. Each mouse was then placed individually into a small compartment (6 × 12 cm) over a Von Frey mesh grid (Bioseb, France) for an additional 45 min session of habituation. Then, von Frey filaments (0.07, 0.6, and 2 g; Bioseb BIO-VF-M) were applied five times on each hind paw and the withdrawal response was scored. The percentage of response was averaged for each filament and compared between genotypes. We also applied a soft brush on the paw to assess responsivity to non-innocuous light touch. Furthermore, we assessed the mechanical threshold response of each mouse using the up-and-down method [23].

### Spontaneous response to skin deformation

The protocol was developed to study the behavioral response to a localized skin deformation and was inspired by Shrestha and Stoeber report describing in details the skin deformation following intradermal injections (PMID:30213982). The day before the injections, the neck of the mice was shaved with an electric clipper. A single intradermal injection of 50 µl of NaCl (0.9%) in the nape of the neck was applied and the subsequent behavioral response was recorded. This type of injection is known to apply positive strain on the dermis and the epidermis, creating skin deformation, or skin wheal (PMID: 30213982). The video was then analyzed and the grooming, scratching, and digging responses were scored by an experimenter (BORIS v8.24.1), blind to the genotypes.

### Mechanical itch (i.e. Alloknesis)

The protocol was adapted from an alloknesis protocol validated in mice [24]. The fur of the nape of the neck was shaved one day before testing and mice were habituated to Plexiglas boxes for a minimum of 3 days prior to testing. The alloknesis response was assessed on a separate testing day. Each mouse received five innocuous mechanical stimuli from three Von Frey filaments (0.02, 0.07 and 0.4 g; Bioseb BIO-VF-M). The scratching response following each stimulation was scored and the scratching percentage was analyzed for each filament.

### Manual analysis of grooming and scratching behaviors

Mice were placed in arenas containing a thin layer of bedding and were video recorded for 20 min (Fig. 1e,f) or, during alloknesis testing, mice were video recorded before and after subcutaneous injections for 20 min (Fig. 1g,h, Fig. 3c,d and Fig. 3 f,g). Videos were analyzed offline (BORIS v8.24.1 [25]) by a blind experimenter, who scored the grooming (i.e. using forelimbs to groom or licking body parts) and scratching behaviors (i.e. using hindlimbs to scratch the neck area).

**Fig. 1.**
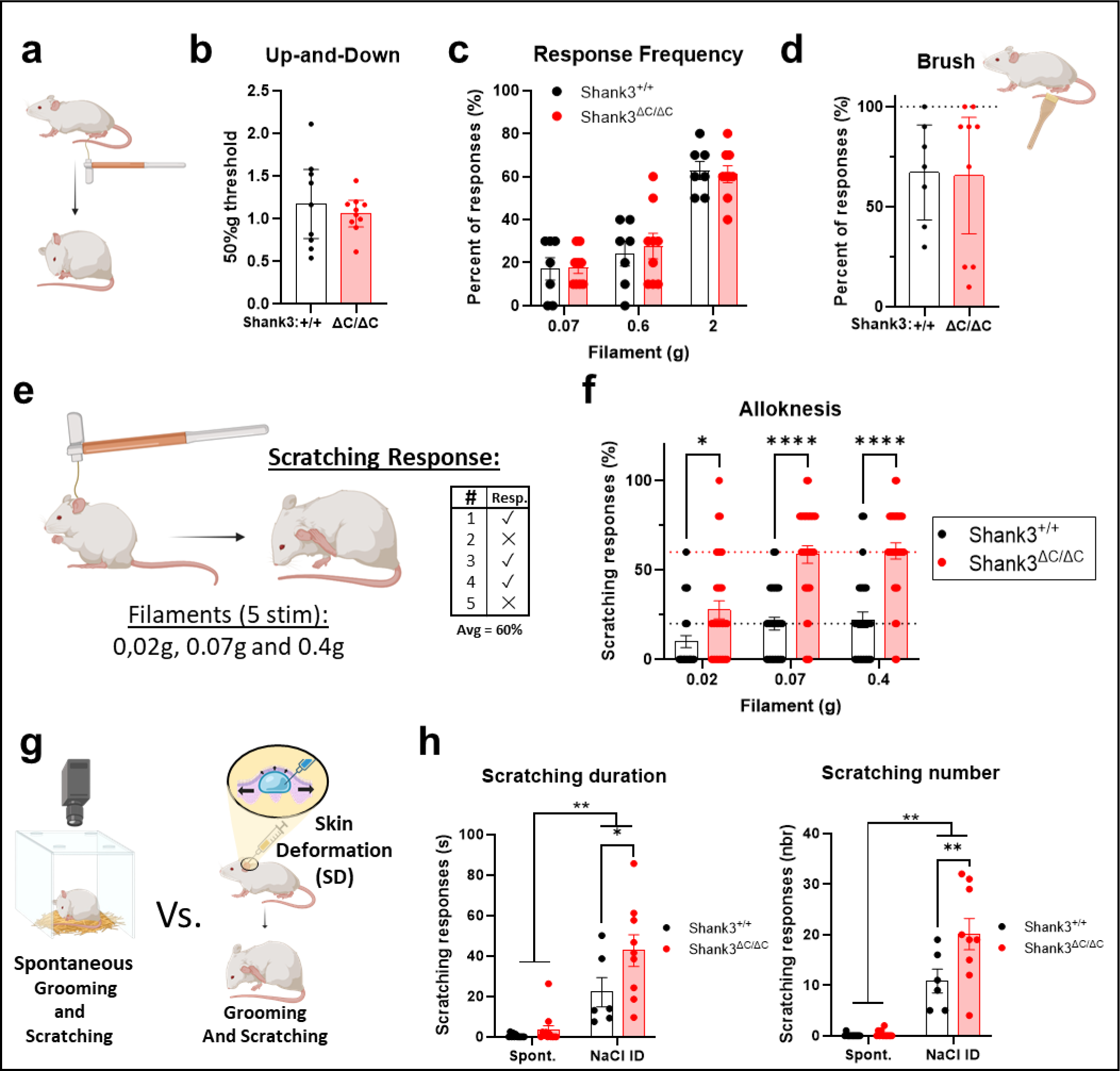
Shank3^ΔC/ΔC^ mice display hypersensitivity to mechanical itch on hairy skin. **a.** illustration of the Von Frey filament experiment, with calibrated filaments applied to the paw of the mice, and the behavioral responses are inspected. **b.** With the up-and-down method to assess withdrawal thresholds, no difference was observed between the groups: t(17) = 0.62, p = 0.5435, n =9 Shank3^+/+^ /10 Shank3^ΔC/ΔC^. **c.** No differences were visible in the frequency of responses to three different filaments (0.07, 0.6 and 2 g); Filament F(1.691, 23.68) = 53.5, p < 0.0001, Genotype F(1, 14) = 0.0365, p = 0.8513, FxG interaction F(2,28) = 0.165, p = 0.8488, n =7 Shank3^+/+^ /9 Shank3^ΔC/ΔC^. **d.** The percent of responses to brush strokes was assessed and did not differ between Shank3^+/+^ and Shank3^ΔC/ΔC^ mice: t(14) = 0.095, p = 0.5435, n =7 Shank3^+/+^ /9 Shank3^ΔC/ΔC^. **e.** Illustration of the mechanical itch protocol, with three filaments of different forces (0.02, 0.07 and 0.4 g) applied to the nape of the neck and the scratching response following the stimulation is observed. **f.** The percent of scratching response was higher in Shank3^ΔC/ΔC^ animals compared to Shank3^+/+^: Filament F(1.794, 102.3) = 23.39, p < 0.0001, Genotype F(1,57) = 42.79, p < 0.0001. FxG Interaction F(2,114) = 6.17, p = 0.0029; with p = 0.0151 for 0.02 g, p < 0.0001 for 0.07 g and p < 0.0001 for 0.4 g, n=28 Shank3^+/+^ and n=31 Shank3^ΔC/ΔC^. **g.** We compared the scratching behavior in two conditions, when mice were video-recorded for 20 min in a novel environment and following a skin deformation (SD) induced by intradermal injection of NaCl. **h.** We measured the total duration of scratching behaviors and observed a net increase in scratching behavior following the skin deformation (SD) protocol, with more scratching in Shank3^ΔC/ΔC^ mice. Treatment F(1, 35) = 43.7, p < 0.0001, Genotype F(1, 35) = 6.48, p = 0.0155, TxG interaction F(1,35) = 3.67, p = 0.064. Shank3^+/+^ vs. Shank3^ΔC/ΔC^: p = 0.85 for Spontaneous (Spont.) and p = 0.016 for SD. We quantified the number of scratching bouts and also observed a higher number of scratching bouts in Shank3^ΔC/ΔC^ mice following skin deformation (SD). Treatment F(1, 35) = 84, p < 0.0001, Genotype F(1, 35) = 8.28, p = 0.007, TxG interaction F(1,35) = 7.43, p = 0.01. Shank3^+/+^ vs. Shank3^ΔC/ΔC^: p = 0.899 for Spontaneous (Spont.) and p = 0.002 for SD, n=6 Shank3^+/+^ and n=9 Shank3^ΔC/ΔC^. Unpaired t-tests were applied in panels **a.** and **d.** 2-way ANOVAs with repeated measures and Sidak multiple comparison tests were applied for data in panels **c**, **f, and h**. Error bars report SEM.

### Pharmacological injections before mechanical itch testing

After mechanical baseline testing, mice were attributed to experimental groups for pharmacological manipulations. On the day of the test, mice were first acclimatized to the test room, they were then placed 30 minutes in the mechanical itch testing chambers for habituation. Following this, they received intradermal injections, and the injection site was marked. Finally their scratching responses to mechanical stimuli were evaluated 45 min post-injection, with the 0.07 and 0.4 g VonFrey filaments. Drugs were prepared in 0.9% NaCl solution. Lidocaine (5 %) combined with QX-314 (0.2 %) was used, as it was shown to be an effective way to induce local anesthesia [26]. Flagellin (0.3 µg, Flg) was administered with QX-314 (0.2 %) as previously described [27]. TAFA4 (200 µg/ml) was administered subcutaneously as previously described [28] or intradermally. Intradermal injections were performed with 31 G syringes and a constant volume of 50 µl was injected. Before injection, each mouse was restrained and the injection site was labeled with a permanent marker to localize the area for filament application.

### Application of gentle touch (GT)

Mice were trained for 10 days, over 2 weeks, in a novel environment (a housing cage 391 x 199 x 160 mm) with a soft brush (SAVITA, facial brush, Amazon). Gentle touch (GT) stroking was performed by an experimenter who was blind to mice genotypes. The hairy back skin of mice was gently stroked with the brush from the nape of the neck to the lumbar part at a constant speed as previously described [29]. The stroking was performed for 10 sessions with three trials within each session (one trial included 100 s of stroking at a rate of 1 stroke per second, and 5 min delay between two trials). Animals were not restrained during the application of the stroking.

### GT as an external procedure before alloknesis assessment procedure

On the testing day, mice were habituated to the apparatus and then tested subsequently with 2 filaments (0.07 and 0.4 g). The test was composed of three steps: First, a baseline response was assessed for each filament to measure the level of alloknesis. Then each animal received 100 brush stimulations. Finally, alloknesis level was tested again in response to the two different filaments (Fig. 4a). There was a delay of 30 min between the baseline session and the post-brushing session.

### GT after the skin deformation procedure

Mice were habituated to the apparatus and then received one intradermal injection in order to induce localized skin deformation. We, then, applied 100 brush stimulations before recording and analyzing the scratching responses (Fig. 4d).

### Real Time quantitative PCR (RT-qPCR)

To evaluate the influence of the deletion of SHANK3 on TRPV1, TRPA1, PIEZO2, TACAN, HisR3 and TAFA4 mRNA levels, total RNA from mice DRGs was extracted according the manufacturer’s instructions of the Nucleospin RNAplus kit (Macherey Nagel, Düren, Germany). Then 500 ng of total RNA was reverse transcribed with PrimeScript™ RT reagent Kit (Takara Bio). For quantitative PCR, 2 ng of cDNA was added as template to SyberGreen master mix (Roche, Basel, Switzerland) and 250 nM primers (Sigma-Aldrich). RT-qPCR was performed on LightCycler® 480 real-time PCR system (Roche). Experiments were performed in triplicates for each experimental group. Relative quantification was achieved according to the comparative method 2^-ΔΔCt^ with HPRT and GAPDH as reference genes [30]. The results are presented as relative expression levels of genes of interest in Shank3^ΔC/ΔC^ mice versus those obtained from control mice. When appropriate, unpaired t-tests or Mann-whitney U tests were applied to compare the mean value for Shank3^ΔC/ΔC^ vs. Shank3^+/+^ mice. The experimenters who performed the dissection, reactions, and analysis were blind to the animal genotypes during the experiments. The assignment of the genotype to the samples was done by the experimenters or colleagues after analysis.

### Electrophysiological properties of dorsal nerves

The protocol was adapted from [31]. The dorsal thoracic nerves, between T6 and T10, were dissected and placed in a three compartments recording chamber filled with artificial cerebrospinal fluid (ACSF), which contained (in mM) 126 NaCl, 3 KCl, 2 CaCl_2_, 2 MgSO_4_, 1.25 NaH_2_PO_4_, 26 NaHCO_3_ and 10 dextrose, pH 7.4–7.5. The extremities of each nerve were placed into a compartment and were separated by a 1 cm long central compartment (illustrated in Supplementary Fig. 3a). The nerves were continuously perfused with warm ACSF (35-36 °C) at a flow rate of 1-2 ml/min. The distal end was stimulated supramaximally (40 µs duration) through two electrodes insulated with Vaseline, and recordings were performed at the proximal end. Signals were amplified, digitized at 500 kHz and recorded. The maximal conduction velocity (CVmax) was measured for each subset of sensory fibers (Aβ-CV > 10 m/s, Aδ-1.2 < CV < 10 m/s or C-fibers CV < 1.2 m/s) and compared between genotypes. Male mice around P100-140 were used (n = 4 mice per genotype).

### *ex vivo* skin-nerve recordings

The isolated dorsal hairy skin-nerve preparation and single-fiber recording technique were used and adapted from previous reports [32–35]. The skin was placed with the corium side facing up in the cell chamber filled with synthetic interstitial fluid (SIF) consisting of (in mM): 120 NaCl, 3.5 KCl, 0.7 MgSO4, 01.7 NaH2PO4, 5 Na2HCO3, 2 CaCl2, 9.5 Na-gluconate, 5.5 glucose, 7.5 sucrose and 10 HEPES at pH 7.4. Fibers in the skin were stimulated with a mechanical search stimulus applied with a custom made probe and were classified by conduction velocity (C-fibers with CV < 1.2 m.s^-1^, Aδ-with 1.2 < CV < 10 m.s^-1^, and Aβ-with CV > 10 m.s^-1^) following a short electric stimulation applied on the receptive field, as previously described [33, 36, 37]. For each identified fibers, a repetitive mechanical stimulation protocol was applied with calibrated ramp and hold stimuli of 5 s that was applied on the receptive field with a cylindrical probe (approximately 0.5 mm of diameter), and which was mounted on a micromanipulator (Physik Instrumente, Karlsruhe, Germany). The actuator (V-273 VC Linear Actuator) vertical displacement was increased by 100 μm each step at the speed of 1000 μm/s (controlled by pClamp software). The exact force applied to the skin surface was measured by the force sensor at the tip of the actuator holding the mechanical probe (V-273.441). Extracellular potentials were recorded with a DAM-80 AC differential amplifier (WPI) and pClamp software (Molecular devices). Offline analysis was performed with Spike2 software (Cambridge Electronic Design) and spikes were identified and analyzed with the principal component analysis extension of the Spike2 software (WaveMark analysis). The experiments and the analysis of individual fiber recordings were carried out under blind conditions until pooling all the results using tattooed markings used for genotyping.

### Statistics

All statistics were performed with GraphPad Prism (Versions 8 and 10). The ROUT (Q = 1%) method was used to identify and remove outliers. The normality of sample distributions was assessed with the Shapiro–Wilk criterion and when violated non-parametric tests were used. When normally distributed, the data were analyzed with unpaired t tests, 1-way ANOVA or repeated measures 2-way ANOVA when appropriate. When non-normal, data was analyzed with Mann–Whitney U tests. For ANOVAs normality of sample distribution was assumed, and followed by Sidak post hoc test. Data are represented as the mean ± SEM and the significance level was 5 %.

### Use of Large Language Models (LLMs) for manuscript preparation

During the preparation of this work Damien Huzard used ChatGPT 4.0 in order to correct and improve English writing solely. After using this tool, DH, AF and JD reviewed and edited the content as needed. DH takes full responsibility for the content of the publication.

## Results

### Shank3^ΔC/ΔC^ mice display hypersensitivity to mechanical itch on hairy skin

Before investigating the impact of the Shank3^ΔC/ΔC^ mutation on somatosensory processing, we first validated the behavioral phenotype of this mouse model in our experimental conditions using protocols classically used to investigate ASD mouse models [38, 39]. Overall, we found that in comparison to the control Shank3^+/+^ animals, Shank3^ΔC/ΔC^ mice did not display social preference (Supplementary Fig. 1a), buried less marbles (Supplementary Fig. 1b), expressed more repetitive self-grooming behavior (Supplementary Fig. 1c), dug less the cage bedding (Supplementary Fig. 1d), traveled less distance during habituation to an experimental cage or during the marble burying testing (Supplementary Fig. 1e), and displayed less burrowing behavior (Supplementary Fig. 1f). All these phenotypes are well validated behavioral alterations in preclinical models of ASD [40–42].

Excessive self-grooming behavior can be associated with repetitive behavior which is a main symptom of ASD [1]. This symptom is believed to have a central origin in ASD mouse models. However, grooming may also be triggered by a peripheral somatosensory hypersensitivity which has never been tested. To explore the somatosensory reactivity of Shank3^ΔC/ΔC^ animals we tested their reactivity to static mechanical stimulations using Von Frey filaments ²(Fig. 1a). Using the frequency and the up-and-down methods [23], there were no differences in hind paw mechanical sensitivity between both genotypes (Fig. 1b and 1c). We also explored sensitivity to dynamic light mechanical stimulation, by applying a gentle brushing on the paw, but did not observe any difference (Fig. 1d).

Then, to assess the mechanical low threshold sensitivity on hairy skin, we tested the Shank3^ΔC/ΔC^ mouse model in a mechanical itch protocol [24, 43]. Using three Von Frey filaments of different forces (0.02, 0.07, and 0.4 g), which were manually applied to the nape of the neck, we assessed the percentage of scratching responses (Fig. 1e). Strikingly, we observed that scratching induced by light mechanical stimuli was more important in Shank3^ΔC/ΔC^ mice compared to Shank3^+/+^ animals (Fig. 1f, Supplementary Fig. 1k). This difference was mild but already significant with the lightest filament and was obvious for the two heaviest filaments, with Shank3^+/+^ mice scratching in average 22.14 % (± 4.41) of the time whereas Shank3^ΔC/ΔC^ mice responded to 60.64 % (± 4.49) of stimulations ().

Additionally, we investigated the spontaneous behavioral response (scratching and self-grooming) to a novel environment in comparison to a localized light skin deformation and mechanical challenge [44], induced by an intradermal saline injection (Fig. 1g). We observed almost no spontaneous scratching behavior in response to a novel environment in both groups (Fig. 1h). However, there was an important increase in scratching duration and bouts (Fig. 1h) in response to a localized skin deformation in both genotypes. This latter was significantly more pronounced in Shank3^ΔC/ΔC^ compared to Shank3^+/+^ mice. Indeed, following this cutaneous mechanical challenge, Shank3^ΔC/ΔC^ mice scratched 20.1 times (± 3.1) for a total duration of 42.7 s (± 7.9) in comparison to Shank3^+/+^ animals, who scratched 10.8 times (± 2.3) for a total duration of 22.1 s (± 7.3). There was a higher spontaneous amount of self-grooming behaviors in Shank3^ΔC/ΔC^ mice in comparison to Shank3^+/+^ animals (Supplementary Fig. 1h) but no difference in response to skin deformation (Supplementary Fig. 1j). These data suggest an overall hypersensitivity to mechanical challenge at the level of the hairy skin in the Shank3^ΔC/ΔC^ mouse model of ASD.

Since the mechanical hypersensitivity to light touch was observable in the nape of the neck, but not on the paw, we decided to focus the screening of the somatosensory alterations at the level of the back skin. Thus, the responsiveness properties of the nerves innervating the back skin of control and Shank3^ΔC/ΔC^ mice were investigated. Skin-nerve recordings were first used to measure the electrophysiological properties of the somatosensory neurons responding to light touch, namely the low-threshold-mechanoreceptors (LTMRs).

### C-LTMRs of Shank3^ΔC/ΔC^ mice have deficits in electrophysiological properties

First, the electrophysiological properties of the entire nerves innervating the hairy skin of the back of the mice were analyzed and the nerve conduction velocity of the different sensory nerve populations was examined (Supplementary Fig. 3a). Based on the maximum conduction velocity (CVmax) of different nerve potential waves, the gross populations of Aβ-, Aδ-or C-fibers could be readily inferred (as illustrated in Supplementary Fig. 3a). There were no differences in the CV of C-, Aδ- and Aβ-fiber populations (Supplementary Fig. 3b).

Then, we aimed to specifically analyze the electrophysiological properties of the mechanosensory fibers. For that, an *ex vivo* skin-nerve preparation was used, as previously described [35, 45], to determine the mechanosensitivity of the different somatosensory fibers innervating the back skin of our mouse model (Fig. 2a). Since the behavioral data described differences in alloknesis to very light filaments, we focused our screening to fibers responding to low mechanical forces (Aβ-, Aδ- and C-LTMRs).

**Fig. 2.**
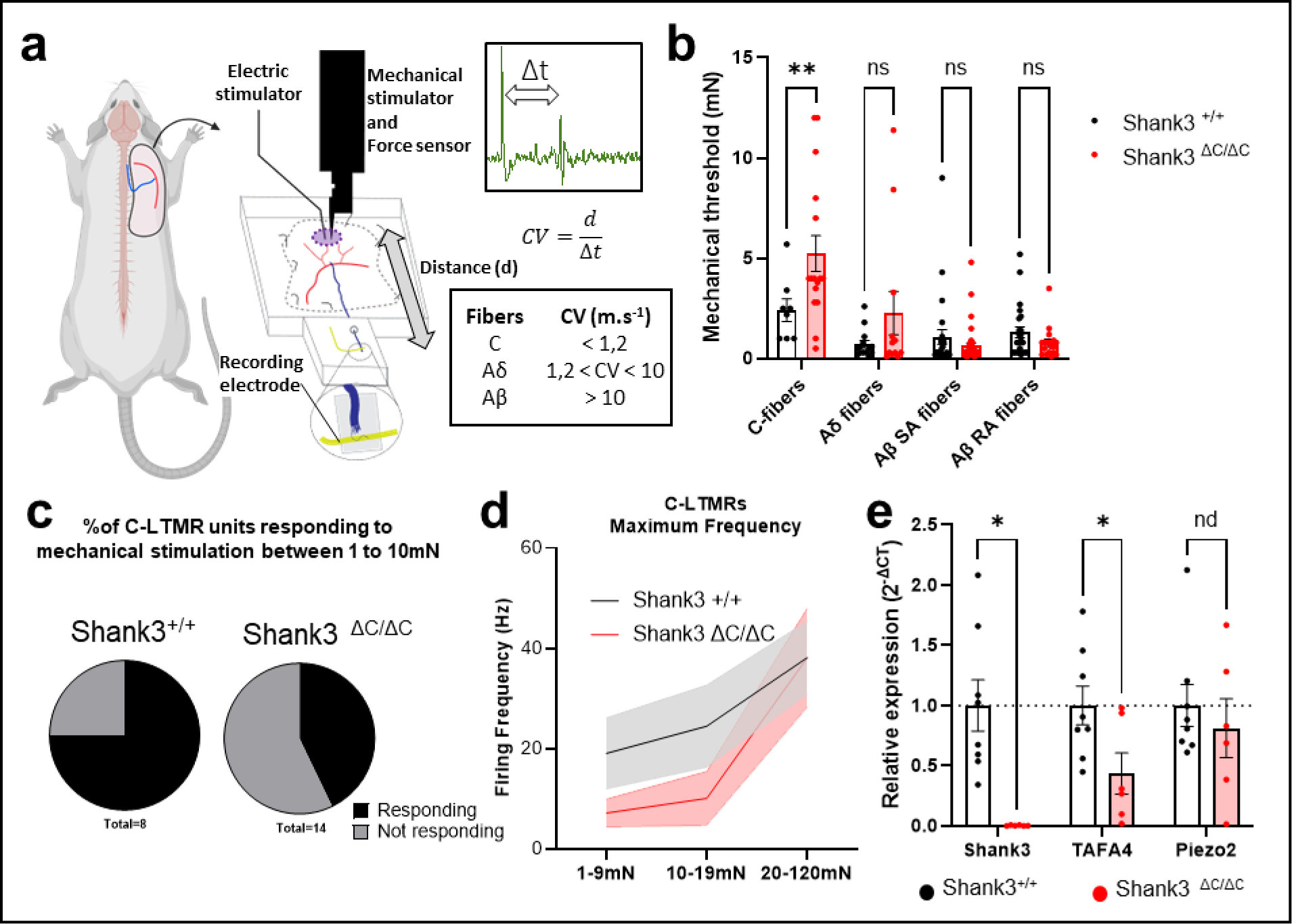
Shank3^ΔC/ΔC^ C-LTMRs are hypofunctioning. **a.** Illustration of the skin-nerve recording procedure performed on the nerves for the back skin on the mice (between T6-T10). Each fiber type was classified based on its CV, and we assessed the minimal mechanical threshold evoking an electrophysiological response (**b, c, d**). **b.** Mechanical threshold of LTMR fibers. 2-way ANOVA with Sidak multiple comparison tests, Fiber type F (3, 150) = 13,86, p < 0.0001, Genotype F (1, 150) = 6,486, p = 0.0119. FxG Interaction F(3,150) = 5.518, p = 0.0013. Shank3+/+ vs. Shank3ΔC/ΔC: Adjusted p value = 0.0030 for C-LTMRs. **c.** Percentage of C-LTMR units responding to mechanical stimulation between 1 and 10mN. **d.** There is no difference in C-LTMRs maximum firing frequency in reponse to increasing forces above 10mN. 2-way ANOVA. **e.** Relative expression of Shank3, TAFA4, and Piezo2 in DRG of Shank3+/+ vs. Shank3ΔC/ΔC mice, assessed by RT-qPCR. 2-way ANOVA with Two-stage linear step-up procedure of Benjamini, Krieger and Yekutieli, Fiber type Gene F (1,874, 22,48) = 4,897, p = 0.0188, Genotype F (1, 12) = 7,471, p = 0.0182. GxG interaction F (2, 24) = 4,897, p = 0.0165. Shank3+/+ vs. Shank3ΔC/ΔC: individual p value = 0.0022 for Shank3 and 0.0347 for TAFA4. Error bars report SEM.

We observed that C-LTMRs of Shank3^ΔC/ΔC^ mice presented a decrease in mechanical sensitivity resulting in a higher mechanical threshold (5.23 mN ± 0.89 for Shank3^ΔC/ΔC^ and 2.41 mN ± 0.56 for Shank3^+/+^; Fig. 2b). Furthermore, there were overall less C-LTMRs responding to low mechanical forces (below 10mN) in Shank3^ΔC/ΔC^ mice compared to WT mice (75% vs 43% respectivly, Fig.2 c). Above 10mN, C-LTMRs respond similarly to increasing forces (Fig. 2d). However, the conduction velocity of C-LTMRs was similar in Shank3^ΔC/ΔC^ and WT mice (Supplementary Fig. 3d). We analyzed similar parameters for Aδ- and Aβ-LTMRs (analyzed as slowly-adapting (SA) or rapidly-adapting (RA) fibers). There were no differences in both mechanical thresholds of Aδ- or Aβ-LTMRs (Fig. 2b) nor differences in their conduction velocities (Supplementary Fig. 3d).

These results illustrate that, in Shank3^ΔC/ΔC^ mice, among all the LTMRs, only a hyporesponsivity of the C-LTMRs could be related to their hypersensitivity to mechanical challenges in the hairy skin.

### Shank3^ΔC/ΔC^ mice have lower expression of TAFA4 in the DRGs

To understand the mechanisms responsible for the mechanical hypersensitivity phenotype in the Shank3 mouse model, we analyzed gene expression from DRG neurons of Shank3^ΔC/ΔC^ and WT mice using RT-qPCR. We targeted our analysis toward gene encoding for proteins associated with C-LTMRs and touch detection, such as TAFA4 and Piezo2. There was a differential expression of the Shank3 gene, with its absence in Shank3^ΔC/ΔC^ mice (Fig. 2e), validating the effectiveness of the genetic mutation. Further analysis revealed a reduced expression of the chemokine TAFA4 in Shank3^ΔC/ΔC^ mice (Fig 2e), suggesting a potential role of Shank3 in regulating TAFA4 expression. Interestingly, no significant differences were observed in the transcript levels of other somatosensory-related genes such as Piezo2, or Trp channels (Supplementary Fig. 3e). We also verified that DRG population proportions were not altered and especially if C-LTMRs were over- or under-represented in thoracic DRGs (Supplementary Fig. 4), which is not case. Combined with the *ex vivo* electrophysiological data, these results point towards an altered responsiveness of C-LTMRs in shank3^ΔC/ΔC^ mice.

### Pharmacological manipulation of Aβ-, but not C-LTMRs abolished the alloknesis hypersensitivity

Based on the electrophysiological and transcriptomic experiments, we wanted to investigate whether C-LTMRs are involved in the pathological alloknesis responses observed in the shank3 mutant mice. Intradermal injections of various pharmacological compounds were performed in the skin of the neck to manipulate specific sensory fibers, known to be involved in alloknesis, before observing the scratching responses to the drug and testing the alloknesis response following the treatment (Fig. 3a).

**Fig. 3.**
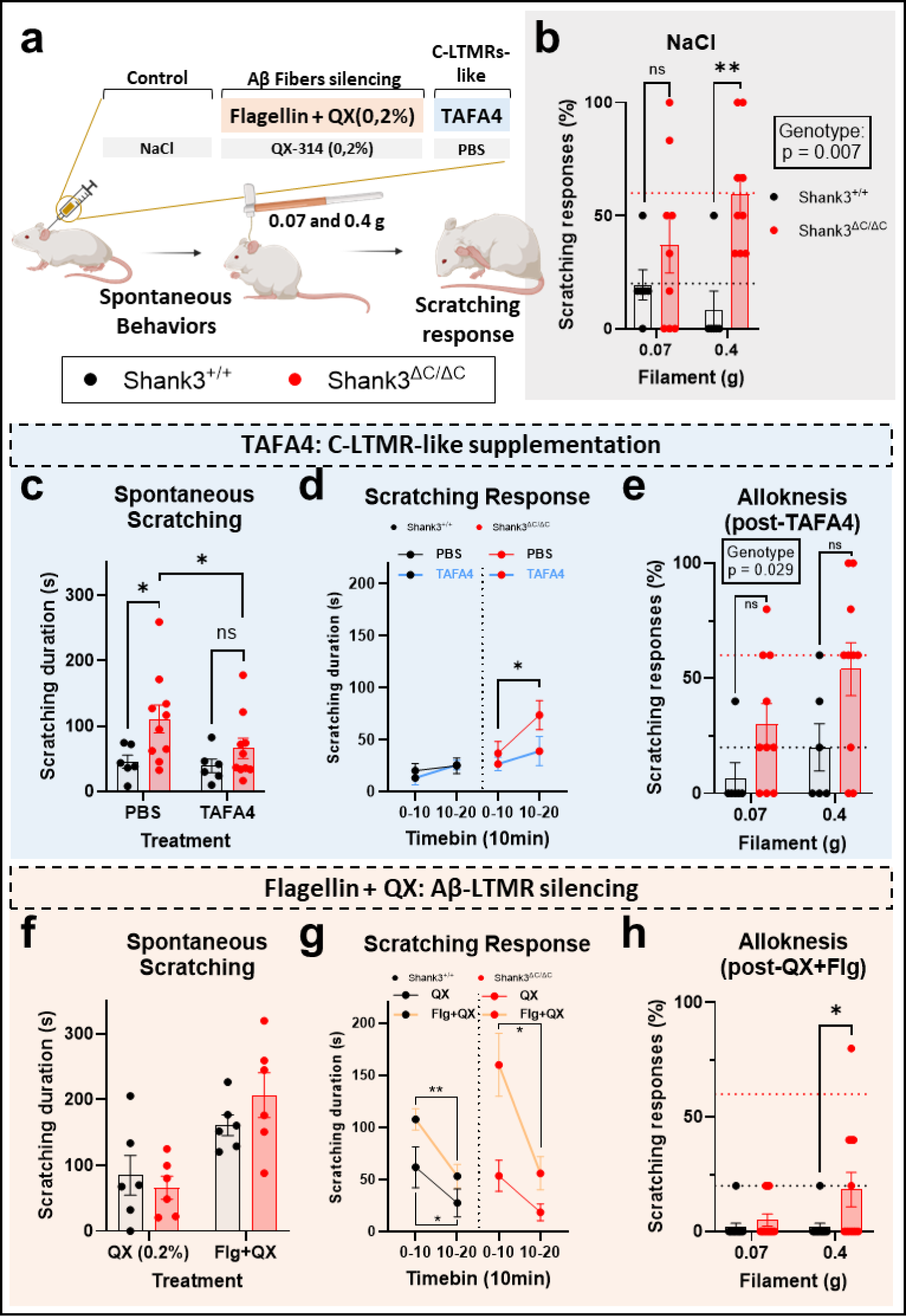
Pharmacological manipulation of Aβ-, but not C-LTMRs abolished the alloknesis hypersensitivity. **a.** Illustration of the experimental procedure. Mice are injected intradermally 45 min before the mechanical itch test. **b.** Following NaCl injection Shank3^ΔC/ΔC^ mice had a higher scratching response. Filament F(1,13) = 0.318, p = 0.582, Genotype F(1,13) = 10.03, p = 0.0074. FxG Interaction F(1,13) = 2.865, p = 0.1143. Shank3^+/+^ vs. Shank3^ΔC/ΔC^: p = 0.422 at 0.07 g and p = 0.0036 at 0.4 g. **c.** Shank3^ΔC/ΔC^ animals scratched more than Shank3^+/+^ mice after PBS injection, but not following a TAFA4 intradermal treatment. Treatment F(1,14) = 2.24, p = 0.157, Genotype F(1,14) = 5.73, p = 0.031. TxG Interaction F(1,14) = 1.27, p = 0.279. Shank3^+/+^ vs. Shank3^ΔC/ΔC^: p = 0.017 after PBS and p = 0.305 After TAFA4. No difference between PBS and TAFA4 injection in Shank3^+/+^ mice (p = 0.818) but TAFA4 decreased scratching response in Shank3^ΔC/ΔC^ mice (p = 0.050). **d.** time course of the scratching response to the PBS and TAFA4 treatments for the two genotypes. TAFA4 decreased the scratching response in Shank3^ΔC/ΔC^ mice only (Shank3^ΔC/ΔC^ mice: p = 0.04 following PBS and p = 0.63 following TAFA4; Shank3^+/+^ mice: p = 0.81 following PBS and p = 0.279 following TAFA4). **e.** Following intradermal TAFA4 injection Shank3^ΔC/ΔC^ mice had a higher scratching response. Filament F(1,14) = 3.898, p = 0.0684, Genotype F(1,14) = 5.915, p = 0.029. FxG Interaction F(1,14) = 0.3182, p = 0.5816. Shank3^+/+^ vs. Shank3^ΔC/ΔC^: p = 0.134 at 0.07 g and p = 0.0639 at 0.4 g. **f.** No difference in scratching between Shank3^+/+^ and Shank3^ΔC/ΔC^ animals after QX-314 injection and important increase in scratching behavior for both genotypes following treatment with Flagelin + QX-314. Treatment F(1,10) = 41.2, p < 0.0001, Genotype F(1,10) = 0.17, p = 0.685. TxG Interaction F(1,10) = 3.64, p = 0.085. Shank3^+/+^ vs. Shank3^ΔC/ΔC^: p = 0.847 after QX-314 and p = 0.396 after Flagellin+QX-314 treatment. Important increase of scratching between QX-314 and Flg+QX-314 injection in Shank3^+/+^ mice (p = 0.019) and in Shank3^ΔC/ΔC^ mice (p = 0.0003). **g.** time course of the scratching response to the QX-314 and Flagellin + QX-314 treatments for the two genotypes. Important immediate scratching response and increased scratching behavior with Flagellin + QX-314 for both genotypes. More scratching in the early phase post-injection (0-10 vs. 10-20 min, p < 0.012) and higher scratching response with Flg+QX-314 in comparison to QX-314 only at the 0-10 min bin (Shank3^ΔC/ΔC^ mice: p = 0.003 at 0-10 min and p = 0.45 at 10-20 min; Shank3^+/+^ mice: p = 0.034 at 0-10 min and p = 0.13 at 10-20 min). Shank3^ΔC/ΔC^ mice: p = 0.488 following QX-314 and p = 0.016 following Flg+QX; Shank3^+/+^ mice: p = 0.035 following QX-314 and p = 0.005 following Flg+QX). **h.** Blockade of Aβ-fibers with Flagellin (0.3 µg) + QX-314 (0.2%) decreased the scratching responses in Shank3^+/+^ and Shank3^ΔC/ΔC^ mice. Filament F(1,21) = 2.87, p = 0.1051, Genotype F(1,21) = 4.281, p = 0.0511 FxG Interaction F(1,21) = 2.87, p = 0.1051. Shank3^+/+^ vs. Shank3^ΔC/ΔC^: p = 0.8472 at 0.07 g and p = 0.0211 at 0.4 g. 2-way ANOVA with repeated measures were applied in panels **b**, **c**, **d**, **e**, **f** and **g**, with Sidak multiple comparison tests. Error bars report SEM.

First, NaCl (0.9%) was injected intradermally as a control condition and the scratching responses to two filaments (0.07 and 0.4 g) was quantified. Consistent with the hypersensitivity to mechanical itch observed in untreated mutant mice (Fig. 1f), responses to mechanical stimulation following NaCl injection were larger in Shank3^ΔC/ΔC^ compared to their WT littermates (Fig. 3b; 37 % ± 12.4 and 59.3 % ± 8.8 for Shank3^ΔC/ΔC^ and 19.4 % ± 6.7 and 8.3 % ± 8.3 for Shank3^+/+^).

However, inhibiting all sensory fibers by blocking sodium channels with intradermal injections of lidocaine strongly reduced the scratching responses of both Shank3^+/+^ and Shank3^ΔC/ΔC^ mice (Supplementary Fig. 5b). Solely, a few Shank3^ΔC/ΔC^ mice scratched their neck in response to the 0.4 g filament (17.8 % ± 7). This confirmed that primary sensory neurons mainly drive the excessive mechanical itch in Shank3 mice at the distal end of the skin.

Then we investigated whether the excessive scratching responses in Shank3^ΔC/ΔC^ mice could be dampened by injection of TAFA4, a chemokine-like protein specifically produced by C-LTMRs which has been validated as a potent analgesic agent for pain relief [28, 46]. Following the injection of vehicle (PBS) the spontaneous scratching behavior induced by the injection was more pronounced in Shank3^ΔC/ΔC^ compared to their WT littermates, over 20 min post-injection (Fig. 3c; 110.9 s ± 21.5 for Shank3^ΔC/ΔC^ and 45.4 s ± 10.2 for Shank3^+/+^ and Fig. 3d). Interestingly, this difference was not visible following the injection of TAFA4 (66 s ± 15.6 for Shank3^ΔC/ΔC^and 39 s ± 10.2 for Shank3^+/+^). However intradermal, or subcutaneous, injections of TAFA4 (200 µg/ml) did not alter the hypersensitivity to mechanically induced itch in Shank3^ΔC/ΔC^ or Shank3^+/+^ mice (Fig. 3e and Supplementary Fig. 5d). This indicates an analgesic effect of TAFA4 visible specifically in the scratching response induced by a localized skin deformation in Shank3^ΔC/ΔC^ animals.

To specifically investigate the implication of Aβ rapidly adapting fibers (Aβ-RA) LTMRs, a cell type well described as responsible for the initiation of mechanical itch [43, 47] we took advantage of the membrane-impermeable derivative of lidocaine, QX-314 (0.2 %) combined with flagellin (Flg, 0.3 µg), a TLR5 activator. Injection of both compounds simultaneously allows the entry of QX-314 into TLR5-expressing fibers and the inhibition of sodium channels in these fibers [48]. We first aimed at confirming, as previously suggested [27, 43] that injection of QX-314 was minimally active alone. Notably, there was no difference between Shank3^ΔC/ΔC^ or Shank3^+/+^ mice in scratching duration following the injection of QX-314 (Fig. 3f; 66.1 s ± 17.4 for Shank3^ΔC/ΔC^ and 84.9 s ± 30.2 for Shank3^+/+^ and Fig. 3g). Moreover, there was a slight increase in alloknesis response in Shank3^+/+^ mice but not in Shank3^ΔC/ΔC^ mice following QX-314 in comparison to NaCl injection with the 0.4 g filament stimulation only (Supplementary Fig. 5f; 53.3 % ± 3.5 for Shank3^ΔC/ΔC^ and 41.2 % ± 5 for Shank3^+/+^ mice). The increase in scratching behavior in response to QX-314 alone was transient as illustrated by the diminution in scratching 10 min following the injection (Fig. 3g left panel).

In contrast, intradermal injection of QX-314 (0.2 %) combined with flagellin (0.3 µg) increased the scratching behaviors following injection for both genotypes (Fig. 3f; 206.6 s ± 34.2 for Shank3^ΔC/ΔC^ and 161 s ± 15.6 for Shank3^+/+^), with a higher increase during the early period following injection in Shank3^ΔC/ΔC^ animals. However, 20 min after the injection, spontaneous scratching behavior is drastically reduced in both genotypes (Fig. 3g, right panel). Finally, we assessed the responsiveness to light punctate stimuli 40 min after the injection and observe a complete anti-alloknesis effect with the selective Aβ-LTMR fibers inhibition that is comparable to that resulting from the non-specific effects of lidocaine (Supplementary Fig. 5b).

These results suggest that TLR5 expressing Aβ-LTMRs pathway, known to be the main initiator of mechanically induced itch, is probably not altered in Shank3^ΔC/ΔC^ mice. It also suggests that C-LTMRs may participate in mechanically induced itch inhibition through two different ways, one requiring TAFA4 secretion locally in the skin in response to localized skin deformation, and another TAFA4 independent (as illustrated in Supplementary Fig. 6).

### Gentle stroking of the dorsal hairy skin reduced the mechanical itch response of Shank3dC/dC mice

Finally, we attempted to develop an external and non-invasive approach that could serve as a therapeutic intervention and allow us to reduce the abnormal hypersensitivity of Shank3^ΔC/ΔC^ mice. To do so, we implemented a gentle touch (GT) protocol based on growing research that highlights the significant impact of gentle touch on mice, which includes inducing emotions like pleasure, reward, and positive affective valence [29, 49–51]. After training to receive gentle touch for 2 weeks (3 trials per day of 100 s of stimulations at 1.6 Hz ± 0.2) we tested the mechanical itch response of the mice before and after gentle stroking of their dorsal region (Fig. 4a). While there was no difference in scratching before or after GT in control mice, there was a reduction in alloknesis for Shank3^ΔC/ΔC^ mice in reaction to GT. Indeed, for both filaments (Fig. 4b, 0.07 and 0.4 g), there was an excessive scratching response in Shank3^ΔC/ΔC^ mice in basal condition, with Shank3^ΔC/ΔC^ mice scratching 40 % (± 11.5,) and 53.3 % (± 12) of the time, while control animals never responded to the stimuli. Importantly, there was no scratching difference following the GT application, with Shank3^ΔC/ΔC^ animals scratching only 20 % (± 6.7) and 31.1 % (± 7.5) of the time, while Shank3^+/+^ mice responded to 8 % (± 8) of stimulations for both filaments. The excessive scratching response of Shank3^ΔC/ΔC^ mice was abolished by GT. Additionally, we tested the effect of GT on the scratching response following our localized skin deformation procedure (Fig. 4c). The GT treatment abolished the difference in scratching response between Shank3^ΔC/ΔC^ mice and Shank3^+/+^ mice (Fig. 4d; 65.5 s ± 19.82 for Shank3^ΔC/ΔC^ and 69.9 s ± 18.54 for Shank3^+/+^ after GT) as previously observed without GT. These results illustrate the implication of an alteration of the primary somatosensory system of Shank3^ΔC/ΔC^ mice in their hypersensitivity to mechanical itch. Importantly, it shows that gentle stroking and the somatosensory reaction elicited has potential therapeutic values.

**Fig. 4.**
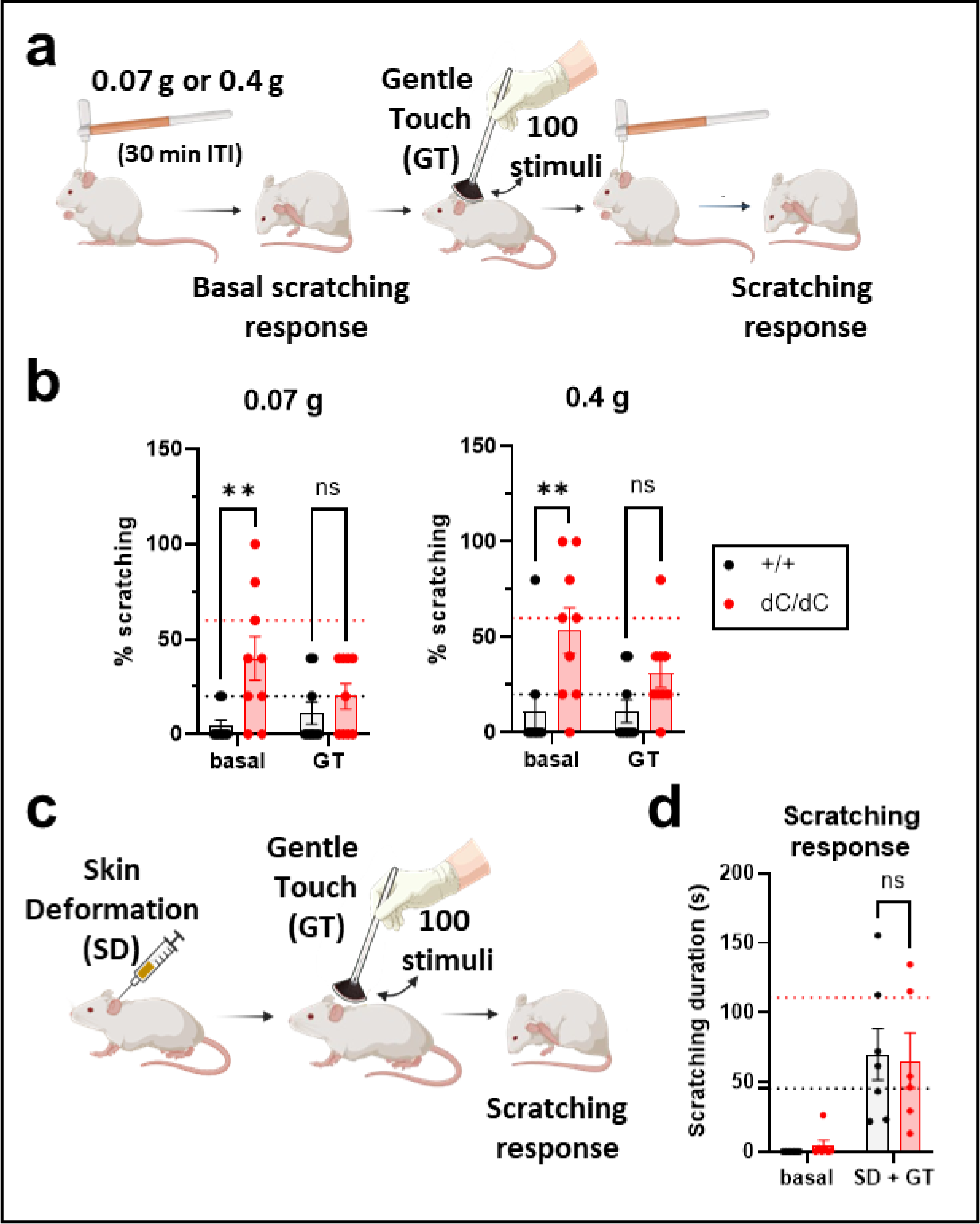
Gentle stroking of the back hairy skin reduced the mechanical itch response of Shank3^ΔC/ΔC^ mice. **a.** Illustration of the experimental procedure. Mice were tested with 2 filaments (0.07 and 0.4 g), with 30 min in-between. Each time, the mechanical itch basal response was assessed, then mice were gently stroked (with 100 stimuli) and the novel mechanical itch response was measured. **b.** There were excessive scratching responses in Shank3^ΔC/ΔC^ mice in basal condition but not following GT when tested with the 0.07 g filament (left, GenotypeF(1,12) = 5.77, p = 0.033, Stroking F(1,12) = 0.55, p = 0.47. GxS Interaction F(1,12) = 3, p = 0.11; Shank3^+/+^ vs. Shank3^ΔC/ΔC^: p = 0.014 at basal and p = 0.62 after GT) and with the 0.4 g filaments (right, GenotypeF(1,12) = 9.59, p = 0.009, Stroking F(1,12) = 0.97, p = 0.34. GxS Interaction F(1,12) = 4.38, p = 0.058; Shank3^+/+^ vs. Shank3^ΔC/ΔC^: p = 0.0028 at basal and p = 0.224 after GT). For Shank3^+/+^ mice there was no effect of the stroking condition, while, for Shank3^ΔC/ΔC^ mice, there was a reduction of scratching following gentle stroking (Filament F(1,8) = 6.13, p = 0.038, Stroking F(1,8) = 6.3, p = 0.036. FxS Interaction F(1,8) = 0.033, p = 0.86. Basal vs. GT: p = 0.096 at 0.07 g and p = 0.065 at 0.4 g). **c.** illustration of the procedure: mice were injected intradermally to induce a localized skin deformation (SD), then 100 GT brush simulations were performed and the behaviors were subsequently analyzed. **d.** Following GT the difference in scratching behavior induced by skin deformation was abolished between Shank3^+/+^ and Shank3^ΔC/ΔC^ mice. Treatment F(1,7) = 17.42, p = 0.004, Genotype F(1,12) = 0.34, p = 0.57. TxG Interaction F(1,7) = 0.13, p = 0.73. Shank3^+/+^ vs. Shank3^ΔC/ΔC^: p = 0.97 at basal and p = 0.8 after GT. 2-way ANOVAs with repeated measures with Sidak multiple comparison tests were applied in panels **b** and **d**. n = 5 Shank3^+/+^ and 8 Shank3^ΔC/ΔC^ mice for panel **b**. n = 3 Shank3^+/+^ and 6 Shank3^ΔC/ΔC^ mice for panel **d** (one Shank3^ΔC/ΔC^ was an outlier after GT (361 s of scratching), statistics remain similar if not included).

## 7. Discussion

The findings of our study highlight several crucial aspects regarding the sensory processing alterations in Shank3^ΔC/ΔC^ mice, a model for ASD, with a specific focus on the scratching behavior elicited by skin deformation or in response to mechanical stimuli. Our observations provide valuable insights into the complex interplay between genetic mutations, neuronal functionality, and behavioral phenotypes in ASD. Indeed, we show, for the first time, that a mouse model of ASD develops a hypersensitivity to itch and higher scratching response to skin deformation. Then, we discovered that these mice, Shank3^ΔC/ΔC^ animals, had an alteration of C-LTMRs’ properties, when previous work on ASD mouse models described a defect of Aβ fibers[17, 18]. Also, gene expression from DRG neurons suggested that the TAFA4 response might also be altered in Shank3^ΔC/ΔC^ mice. Using pharmacological manipulations to specifically target subsets of somatosensory fibers combined with the analysis of spontaneous response to skin deformation and mechanical itch we showed the probable existence of two different pathways, dependent or not on TAFA4, which may be altered in the Shank3^ΔC/ΔC^ mouse model of ASD. Indeed, we showed a potential implication of the Aβ-LTMR fiber in the hyper-expression of alloknesis in Shank3^ΔC/ΔC^ mice as well as a role of TAFA4 in the control of the scratching behavior following skin deformation. Finally, we suggest that gentle touch, activating a large population of C-LTMRs (expressing TH and/or MRGPRB4) could rescue the alloknesis phenotype in Shank3^ΔC/ΔC^ mice as well as their aberrant spontaneous scratching response to skin deformation. Overall, our work paves the way for a new way to study tactile deficits in ASD-related models, with direct implication of peripheral somatosensory deficits related to pathological conditions.

Previous reports using different mouse models of ASD have described hyperreactivity to light touch [17–19], but none have reported a higher susceptibility to itch. Our behavioral assays (Fig. 1) demonstrated that Shank3^ΔC/ΔC^ mice exhibit an increased susceptibility to both localized skin deformation (with an intradermal injection) and mechanical itch, which is illustrated in both situations by an increase in scratching behaviors toward the area stimulated. Our finding aligns with clinical reports suggesting a link between the disturbance of skin sensation and ASD [52]. Moreover, ASD patients have been shown to suffer more from atypical sensory processing [53] and cutaneous alterations [10, 13, 52, 53], such as dermatitis atopic [12]. The abnormal increased sensitivity to mechanical challenges was specifically pronounced in the nape of the neck, but not at the level of the paw, suggesting a potential specific hypersensitivity of hairy skin and not glabrous skin. Our observations confirm that animal models can also be used to study skin hypersensitivity related to ASD. However, It remains to determine whether other ASD mouse models also present itch hypersensitivity in the hairy skin.

The study of mechanical itch has seen significant advancements in recent years, revealing complex neural pathways and mechanisms. The involvement of TLR5+ Aβ-LTMRs in transmitting mechanical itch has been highlighted in recent literature [47, 54]. Interestingly, electrophysiological data suggest that these fast-conducting fibers were not altered in Shank3^ΔC/ΔC^ mice (Fig. 2). However, silencing these fibers was sufficient to reduce alloknesis in this mouse model, suggesting that, independently of the state of other sensory neurons, the mechanical itch response may be excessively initiated by Aβ-RA-LTMRs. It also suggests an interesting potential use as a topical and local therapeutic strategy to reduce mechanically induced itch responses. Another pivotal discovery for the comprehension of Shank3^ΔC/ΔC^ mice somatosensory phenotypes was the role of low-threshold mechanoreceptors, specifically VGLUT3-lineage sensory neurons, which correspond to a subset of C-LTMRs [46, 47] in mediating spinal inhibition of itch by tactile stimuli [55]. Our electrophysiological experiments on back skin (Fig. 2) revealed that, in Shank3^ΔC/ΔC^ mice, electrophysiologically defined C-LTMRs (including multiple genetically identified subpopulation of sensory neurons) are less sensitive to mechanical stimulations. Our qPCR analysis (Fig. 3) also emphasized a decreased expression, in the DRGs, of TAFA4, a key protein associated with VGLUT3+ C-LTMRs [46, 56], in Shank3^ΔC/ΔC^ mice.

Our experiments with TAFA4 injections did not rescue the excessive alloknesis phenotype of Shank3^ΔC/ΔC^ mice but it reduced the behavioral scratching response following a localized skin deformation induced by the injection. This suggests that the alloknesis observed in Shank3^ΔC/ΔC^ might be independent of the C-LTMRs dysfunction or that the C-LTMRs properties contributed to reducing alloknesis are TAFA4-independent. On the other hand, the increased behavioral scratching response of Shank3^ΔC/ΔC^ mice, following skin deformation, could be TAFA4-dependent since we showed that administration of TAFA4 could restore this phenotype in Shank3^ΔC/ΔC^ animals (as illustrated in Supplementary Fig. 6). To note, this disparity between the effect of TAFA4 in rescuing different types of mechanical-induced itch may be the reflection of the diversity of the sensory neurons classified as C-LTMRs; as we know at least two different genetically defined populations of C-LTMRs, those expressing TH and TAFA4, and the other expressing MRGPRB4 [46, 49, 57–59].

Our results obtained using gentle touch stimuli, which are supposed to activate preferentially both populations of C-LTMRs (i.e. expressing TH and/or MRGPRB4), tend to confirm that manipulation of C-LTMRs inputs is key to regulating responses to itch in Shank3^ΔC/ΔC^ animals. Furthermore, this approach suggests a novel and non-invasive therapeutic method to potentially alleviate skin-related symptoms in ASD. Indeed, manual therapeutic interventions are already used in the context of ASD and improving our understanding of the exact pathways involved in the beneficial effect they might provide would greatly improve their application [4].

These results are of importance since C-LTMRs are implicated in the regulation of rewarding and social behaviors in mice [29, 49, 50, 60]. Indeed, social deficits are a hallmark of ASD and its mouse models, and the possibility for the identification of a neuronal target at the skin level has promising therapeutic and clinical implications. Future experiments may investigate the link between C-LTMRs and TAFA4 deficits in the context of sociality in the Shank3^ΔC/ΔC^ mouse model. Moveover, establishing the exact identity of the C-LTMRs subpopulation regulating responses to itch remains to be performed in-depth.

Our study marks a substantial advancement in understanding somatosensory processing in ASD. Unveiling mechanical itch hypersensitivity in a genetic ASD model implies that peripheral sensory changes might be more critical in ASD than earlier assumed. This revelation paves the way for new research directions into ASD’s sensory dimensions and possible therapeutic strategies. Crucially, our results highlight the necessity of an integrated approach to ASD research, encompassing both central and peripheral nervous system elements, to thoroughly unravel the intricate neurobiological basis of the condition. Indeed, future research driven by better clinical assessments is warranted in order to properly understand tactile processing differences in ASD [61].

## Acknowledgements

We would like to thank Julie Perroy for the initial donation of the Shank3 mice and relevant scientific discussions. Jacques Noël for crucial skin nerve recording advice. Marie-Adèle Charldoreille, Oceane Gentilini, Amelyne Marcon David as well as the Biocampus/iexplore team for major help with animal caretaking. Lillian Basso for exciting and relevant scientific exchanges.

## Statement on illustrations

*Illustrations are made to visually help comprehension of the protocols but do not match exact experimental configurations (e.g. mice used in the article had black fur)*.

## Contributions

DH, EB, and AF designed the experiments. DH performed and analyzed the behavioral experiments. MM and CG assisted the mechanical itch experiments. CG performed the *gentle touch* training and the scoring of some behavioral responses. VS performed and analyzed the qPCR experiments. JD and DH performed the nerve recordings, JD performed the electrophysiological analysis, and DH the statistics. DH, GO, and EB performed the skin-nerve recordings and analysis. GG performed some experiments not included in the final manuscript. DH prepared the figures and the first draft of the manuscript. DH and AF wrote the manuscript. EB and JD provided critical feedback and corrections on the final version of the manuscript. All authors validated the final version of the manuscript. EB and AF provided the fundings.

