## Supplemental material and methods for "Primary sensory neuron dysfunction underlying mechanical itch hypersensitivity in a Shank3 mouse model of autism"

#### Supplementary Methods

##### Three-chamber social-preference test

The three-chamber arena was composed of three compartments (20 × 40 × 22 (h) cm each), and separated by two sliding doors (5 × 8 (h) cm). Two stimuli boxes were placed at opposite sides of the apparatus. After room habituation, each animal was placed individually in the arena, starting from the middle compartment, and was let free to explore for 10 min with two empty stimuli boxes. Then, the mouse was locked in the middle compartment and one unfamiliar juvenile mouse (C57B6J/n bred and housed in the IGF facility), was placed in one of the two stimuli box, and an inanimate object (made from Lego blocks, roughly shaped like a mouse) was placed in the other stimuli box. The tested mouse was then allowed to explore the entire apparatus for 10 min. The whole experiment was video-recorded, and animal position was tracked using ANY-maze (Stoelting Europe, Dublin). The side where the stranger mouse and the inanimate object were placed was alternated between each tested mouse to avoid side-bias.

##### 10min cylinder test: Grooming vs. Digging assessment

Mice were placed in a cylinder (diameter = 15 cm approx.) for 10 min. The cylinder contained some bedding (1cm) allowing mice to perform some bedding-displacement behaviors (called "Digging"). All sessions were video-recorded with a camera placed at the top of the cylinders. Videos were then scored by an experimentator using Boris software [1] which analyzed the Digging and Grooming behaviors.

##### Overnight burrowing test

This test was adapted from previous studies [2] and was used to establish the burrowing behavior of the mice illustrating a naturalistic-like capacity. For this test, we built cylinders with two feet on one side (from hydro-alcoholic solution bottles and the feet were made from insulin syringes). The empty cylinders were placed at 9am in the homecage of the mice (isolated or group-housed) and the food dispenser was removed. At 5pm, 200g of food pellets were inserted in the cylinder. The following morning, at 9am, the food left in the cylinder was weighed and the amount of food displaced was calculated.

##### Marble Burying test

As previously described [3] mice were habituated for 2 days to a new cage with 2-3 cm of fresh bedding material. Then, on the test day, 20 marbles were evenly placed following a 4x5 rectangular pattern (using a paper grid with holes for each marble) and left 20 min in the cage. After the 20min, mice were moved back to their homecage and the number of 'marbles untouched' was assessed using the same paper grip and counting the amount of marbles that remained in the exact same spot. A score of 0 was attributed to a hole without a grid inside, a score of 0.5 was attributed to a marble that was partially visible and a score of 1 was counted when the marble was entirely visible inside the hole.

##### Immunohistology

###### *Tissue collection and processing*

Adult mice (P94 to P119) were anesthetized with ketamine and xylazine and then transcardially perfused with phosphate-buffered saline (PBS) followed by 4% paraformaldehyde in PBS. Thoracic DRGs (T8 to T13) were dissected, postfixed in 4% paraformaldehyde for 1 hour, and cryoprotected in 30% sucrose in PBS. Tissues were then frozen in optimum cutting temperature (Tissue-Tek) and sectioned using a cryostat (Leica). DRGs were sectioned at 18 µm on collected on Superfrost Plus Adhesion Microscope Slides (Fisher Scientific) and stored at -20°C

##### *Immunofluorescence*

Tissues were incubated for 1 hour and blocked in a solution consisting of 0.1 M PBS with 0.3% Triton X-100 (Sigma-Aldrich) plus 5% normal donkey serum. Primary and secondary antibodies were diluted in 0.1 M PBS with 0.3% Triton X-100 (Sigma-Aldrich) plus 5% normal donkey serum. Sections were then incubated overnight at RT, under agitation, in primary antibody solution, washed 3 times (5 min for each times) in 0.1 M PBS with 0.3% Triton X-100, incubated for 2 hours in secondary antibody at RT, and washed again 3 times (5 min for each times) in 0.1 M PBS. Sections were then mounted using Dako fluorescence mounting medium. Images were acquired with a Leica SP8 confocal microscope. For all markings, for each mouse, three sections from T11 to T13 DRGs were analyzed and counted. Primary antibodies: The following antibodies were used: anti-TH:Millipore (sheep; 1:500), anti-PGP9.5:Millipore (rabbit;1/500), anti-SP:Neuromics (guinea pig;1/250), anti-NF200:Sigma (mouse;1/200) and anti-CGRP:Immunostar (rabbit;1/100). To identify IB4-binding cells, fluorophore-conjugated IB4-594 (1:500; Vector Laboratories) was used in place of primary and secondary antibodies. Secondary antibodies: Alexa Fluor–conjugated secondary antibodies were acquired from Invitrogen and Jackson ImmunoResearch Labs.

#### Supplementary Figure Legends

**Supplementary Fig. 1 – Additional behavioral Shank3$^{\Delta C/\Delta C}$ phenotypes. a.** Time in the social and non-social zone during the social preference task, Zone F(1,22) = 28.22, p < 0.0001, Genotype F(2,22) = 0.2314, p = 0.7953, ZxG interaction F(2,22) = 2.141, p = 0.141; with Social vs. Empty : p = 0.0011 for Shank3$^{+/+}$ and p = 0.2047 for Shank3$^{\Delta C/\Delta C}$ mice. **b.,** left panel: Number of marbles left undisturbed after the marble burying test, t(53) = 7.185, p < 0.0001. **b.,** right panel: representation of the independent experimental cohorts constituting the results presented in b, left panel. There are no differences between cohorts. Cohort F (1, 51) = 0,3727, p = 0.5443, and genotype F (1, 51) = 49,43, p < 0.0001. **c., d.** during a session of 10 minutes in a neutral cylinder mice were analyzed and the grooming (**c**) and digging (**d**) behaviors were analyzed; **c** t(45) = 3.613, p = 0.0008 and **d** t(45) = 6.209, p < 0.0001. **e.** The distance traveled during the habituation and testing of the marble burying test; Session F(1.986, 73.48) = 43.63, p < 0.0001, Genotype F(1,37) = 19.24, p < 0.0001, SxG interaction F(2,74) = 12.56, p < 0.0001. **f.** Quantity of food left in a cylinder inserted overnight in the homecage with 200g of pellets; t(9) = 6.238, p = 0.0002. **g.** Protocol for spontaneous observation of grooming and scratching behaviors. **h.** the amount of grooming behavior was analyzed (t(22) = 4.629, p = 0.0001). **i.** protocol for the observation of grooming and scratching after skin deformation. **j.** Following skin deformation with a single NaCl intradermal injection in the nape of the neck, we observed that Shank3$^{+/+}$ mice groomed more and reached the same level as Shank3$^{\Delta C/\Delta C}$ animals (132.1 s ± 20.6 for Shank3$^{\Delta C/\Delta C}$ and 108.3 s ± 29.3 for Shank3$^{+/+}$), t(13) = 1.553, p = 0.144. **k.** Representation of the independent experimental cohorts constituting the results presented in Figure 1F. Genotype F (1, 21) = 18,58 , p = 0.0003 for cohort 1, F (1, 13) = 8,825, p = 0.0108 for cohort 2, and F (1, 17) = 16,26 p = 0.0009 for cohort 3. Unpaired t-tests were applied in panels **b left**, **c**, **d, f, h,** and **j**, and 2-way ANOVA was applied for data in panels **a, b right, e,** and **k**, with Sidak multiple comparison tests. Error bars report SEM. Each dot represents one mouse.

**Supplementary Fig. 2 – Shank3$^{\Delta C/\Delta C}$ mice sexual dimorphism in tested behaviors. a.,** left panel: Females Shank3$^{+/+}$ buried less marbles than Shank3$^{+/+}$ males, p = 0.031. **a.,** right panel: representation of the independent experimental cohorts constituting the results presented in a, left panel. There are no differences between cohorts. 3Way ANOVA, corrected for multiple comparisons using Sidak post hoc test. Cohort effect is p = 0.4120, genotype effect is p = 0.0119 and the sex effect is p < 0.0001. **b.** No difference in the amount of grooming between males and females, p = 0.2052. **c** No difference in digging behavior between males and females, p = 0.8024. **d.**, **e.,** and **f.** Mechanical itch testing with no difference according to the sex of the mice. Sex effect is p = 0.7429, 0.336 and 0.4075 for 0.02 g, 0.07 g and 0.4g respectively. Genotype effect is p = 0.145, 0.0006 and p<0.0001 for 0.02 g, 0.07 g and 0.4g respectively. Each dot represents one mouse

**Supplementary Fig. 3 – Shank3$^{\Delta C/\Delta C}$ low threshold mechanoreceptors conduction velocity. a.** Illustration of the procedure to measure overall nerve conduction velocities. Dorsal nerves were dissected out, and mounted in a three-chamber recording system and an electric current was applied to stimulate the different fiber populations (C-, Aδ- and Aβ-fiber populations). The maximal conduction velocity (CVmax) was analyzed. **b.** No difference in the CVmax of C, Aδ, and Aβ-fibers was observed. n = 10 nerves (N = 4 mice) for Shank3+/+ and n = 12 nerves (N = 4 mice) for Shank3ΔC/ΔC animals. **c.** Illustration of the skin-nerve recording procedure performed on the nerves for the back skin on the mice (between T6-T10). Each fiber type was classified based on its responses to low mechanical stimulation (<10mN) and its CV, measured by calculating the latency between an electrical stimulation and the first action potential. **d.** No difference in the CV of C, Aδ, and Aβ-LTMRs was observed. **e.** Additional RT-qPCR analysis on somatosensory-sensory related genes. 2-way ANOVA were applied for data in panels **b, d,** and **e**, with Sidak multiple comparison tests in **b**, and **d** and Two-stage linear step-up procedure of Benjamini, Krieger and Yekutieli in panel **e**. Error bars report SEM. Each dot represents one fiber in **b** and **d**, and one mouse in **e**.

**Supplementary Fig. 4 – No change in TH, SP, IB4, NF200 and CGRP expression in thoracic DRGs in Shank3$^{\Delta C/\Delta C}$ mice. a.** Expression of TH neurons in thoracic DRGs in Shank3$^{+/+}$ (top) and Shank3$^{\Delta C/\Delta C}$ (bottom) mice. **b.** Expression of SP neurons in thoracic DRGs in Shank3$^{+/+}$ (top) and Shank3$^{\Delta C/\Delta C}$ (bottom) mice. **c.** Expression of IB4 neurons in thoracic DRGs in Shank3$^{+/+}$ (top) and Shank3$^{\Delta C/\Delta C}$ (bottom) mice. **d.** Expression of NF200 neurons in thoracic DRGs in Shank3$^{+/+}$ (top) and Shank3$^{\Delta C/\Delta C}$ (bottom) mice. **e.** Expression of CGRP neurons in thoracic DRGs in Shank3$^{+/+}$ (top) and Shank3$^{\Delta C/\Delta C}$ (bottom) mice. **f.** Total number of sensory neurons (corresponding to the green immunostaining). PGP9.5: N=3 animals for Shank3$^{+/+}$ (n = 9 slices per animal) ; N=3 animals for Shank3$^{\Delta C/\Delta C}$ (n = 9 slices per animal). Unpaired t-test was performed for PGP9.5 analysis. **g.** Percentages of neurons expressing TH (corresponding to immunostaining in Supplementary Fig. 3a), SP (corresponding to immunostaining in Fig. 1b), IB4 (corresponding to immunostaining in Supplementary Fig. 3c), NF200 (corresponding to immunostaining in Supplementary Fig. 3d) and CGRP (corresponding to immunostaining in Supplementary Fig. 3e) in Shank3$^{+/+}$ (gray circles) and Shank3$^{\Delta C/\Delta C}$ (red circles) mice. TH, SP, IB4, NF200 and CGRP: N=3 animals for each condition (n = 3 slices per animal). two-way ANOVA was performed for TH, SP, IB4, NF200 and CGRP analyses: Genotype F(1,80) = 0.4544, p = 0.5022, Population F(4,80) = 59,81, p < 0.0001, GxP interaction F(4,80) = 1,442, p = 0,2279. Scale bars, 50 µm. Each dot represents one slice. Slices from the same animals are represented with the same color.

**Supplementary Fig. 5 - Alloknesis responses to additional pharmacological treatments. a** Illustration of the experimental procedure. Mice are injected intradermally 45 min before the mechanical itch test. **b** Following Lidocaine injection there was no more scratching behavior. Filament F(1,13) = 5.363, p = 0.0375, Genotype F(1,13) = 2.511, p =0.1371. FxG Interaction F(1,13) = 2.511, p = 0.1371. Shank3$^{+/+}$ vs. Shank3$^{\Delta C/\Delta C}$: p = 0.999 at 0.07 g and p = 0.0664 at 0.4 g. **c** Shank3$^{\Delta C/\Delta C}$ animals self-groomed more than Shank3$^{+/+}$ mice after PBS injection, but not following a TAFA4 intradermal treatment. Treatment F(1,28) = 0.035, p = 0.852, Genotype F(1,28) = 11.28, p = 0.002. TxG Interaction F(1,28) = 0.75, p = 0.394. Shank3$^{+/+}$ vs. Shank3$^{\Delta C/\Delta C}$: p = 0.006 after PBS and p = 0.089 after TAFA4 treatments. **d** Following subcutaneous TAFA4 injection Shank3$^{\Delta C/\Delta C}$ mice had a higher scratching response. Filament F(1,13) < 0.001 , p = 0.999, Genotype F(1,13) = 24.37, p = 0.0003. FxG Interaction F(1,13) = 0.936, p = 0.429. Shank3$^{+/+}$ vs. Shank3$^{\Delta C/\Delta C}$: p = 0.0052 at 0.07 g and p = 0.0004 at 0.4 g. **e** No difference in self-grooming between Shank3$^{\Delta C/\Delta C}$ and Shank3$^{+/+}$ mice after QX-314 or Flg+QX-314 injections. Treatment F(1,10) = 2.46, p = 0.148, Genotype F(1,10) = 0.116, p = 0.74. TxG Interaction F(1,10) = 0.74, p = 0.41. Shank3$^{+/+}$ vs. Shank3$^{\Delta C/\Delta C}$: p = 0.96 after QX-314 and p = 0.685 after Flg+QX-314 injections. **f** Following QX-314 (0.2%) injection Shank3$^{\Delta C/\Delta C}$ mice had a higher scratching response. Filament F(1,36) = 16.91, p = 0.0002, Genotype F(1,36) = 8.688, p = 0.0056. FxG Interaction F(1,36) = 1.82, p = 0.1857. Shank3$^{+/+}$ vs. Shank3$^{\Delta C/\Delta C}$: p = 0.0048 at 0.07 g and p = 0.2728 at 0.4 g. 2-way ANOVAs with repeated measures with Sidak multiple comparison tests were applied in panels **b**, **c**, **d**, **e** and **f**. Error bars report SEM. Each dot represents one mouse

**Supplementary Fig. 6 - Hypothetical diagram for 2 different pathways involved in the control of Itch and scratching response in Shank3$^{\Delta C/\Delta C}$ mice.** Our experiments identified two distinct pathways for scratching behavior in response to tactile stimuli: punctate mechanical stimulation (defined as direct, external mechanical pressure, highlighted in orange) and skin deformation (characterized by localized mechanical distortion, for example following intradermal injection, highlighted in green). The response to punctate mechanical stimulation is predominantly mediated by A**β**-LTMR fibers expressing TLR5 and is hypersensitive in Shank3$^{\Delta C/\Delta C}$ mice, leading to an alloknesis phenotype (excessive mechanical itch responses). In contrast, skin deformation induces scratching, which can be attenuated by TAFA4 in Shank3$^{\Delta C/\Delta C}$ mice. However, TAFA4 does not reduce alloknesis, suggesting that the skin deformation response is TAFA4-dependent, while the mechanical itch response is TAFA4-independent. Shank3$^{\Delta C/\Delta C}$ mice, with deficient C-LTMRs, exhibit increased susceptibility to itchiness in both scenarios. A supplementation with TAFA4 could only retrieve the spontaneous scratching response to skin deformation in Shank3$^{\Delta C/\Delta C}$ but not their alloknesis hypersensitivity.

### Supplementary Figures

#### Supplementary Fig. 1

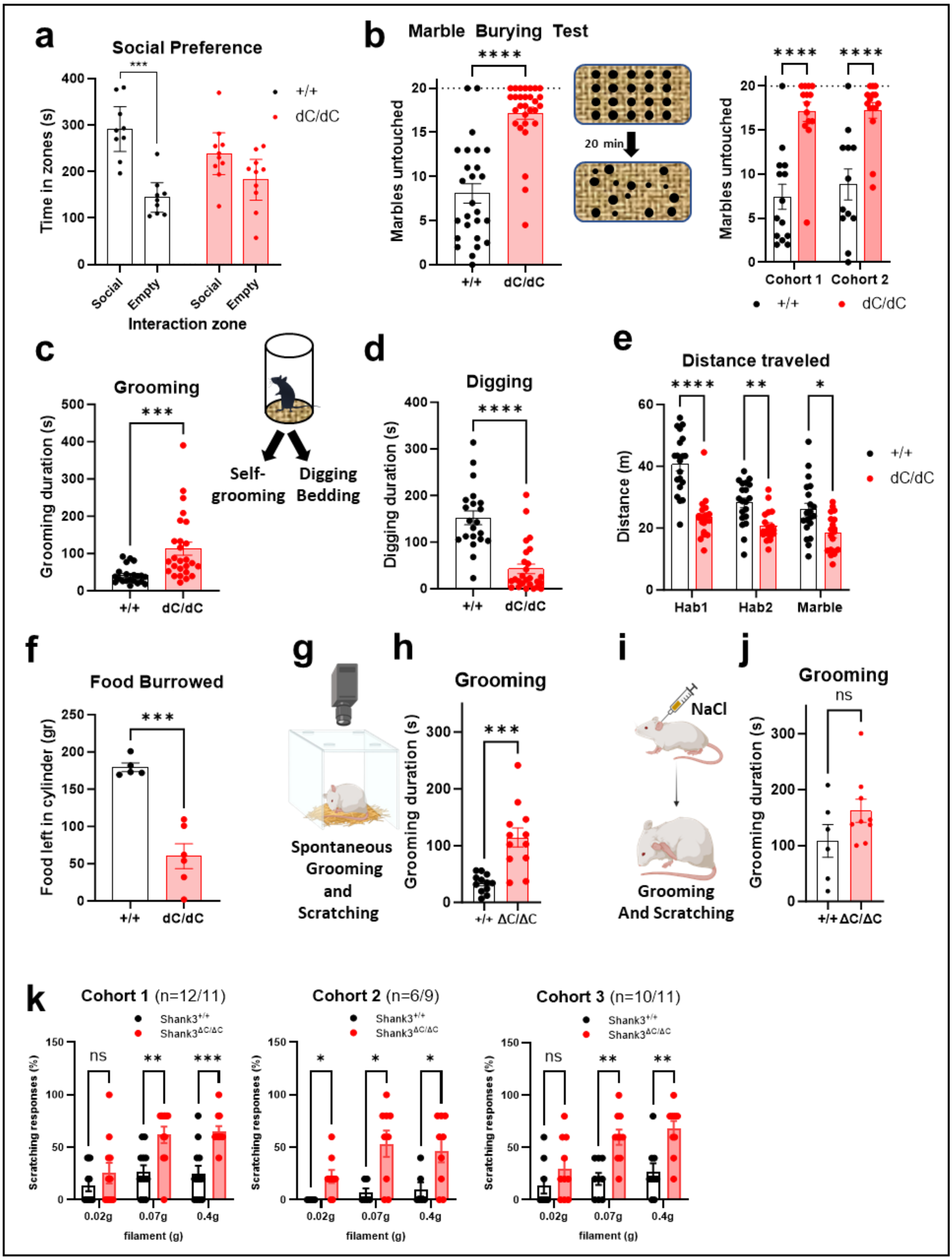

**Supplementary Fig. 2**

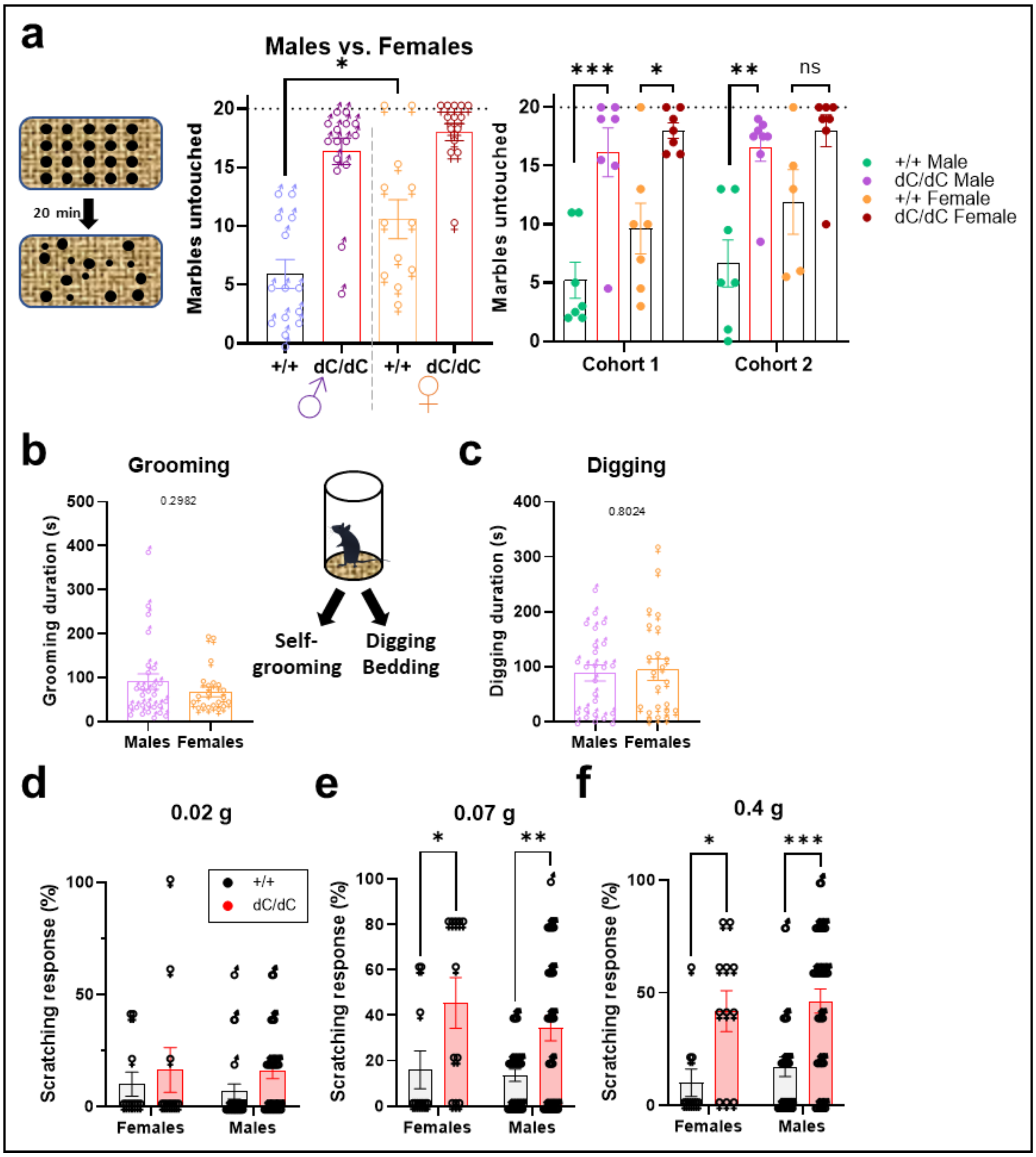

**Supplementary Fig. 3**

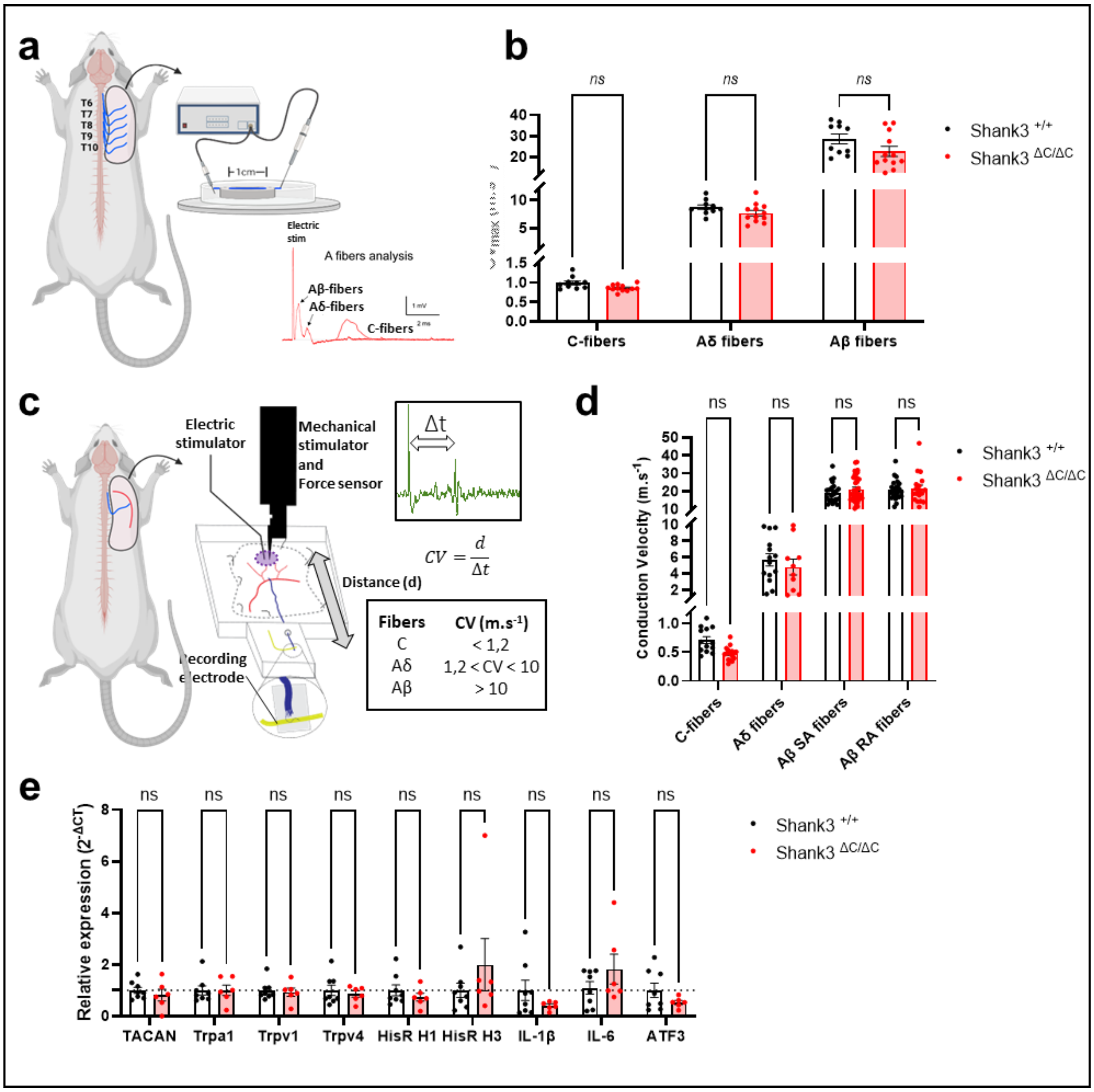

#### Supplementary Fig. 4

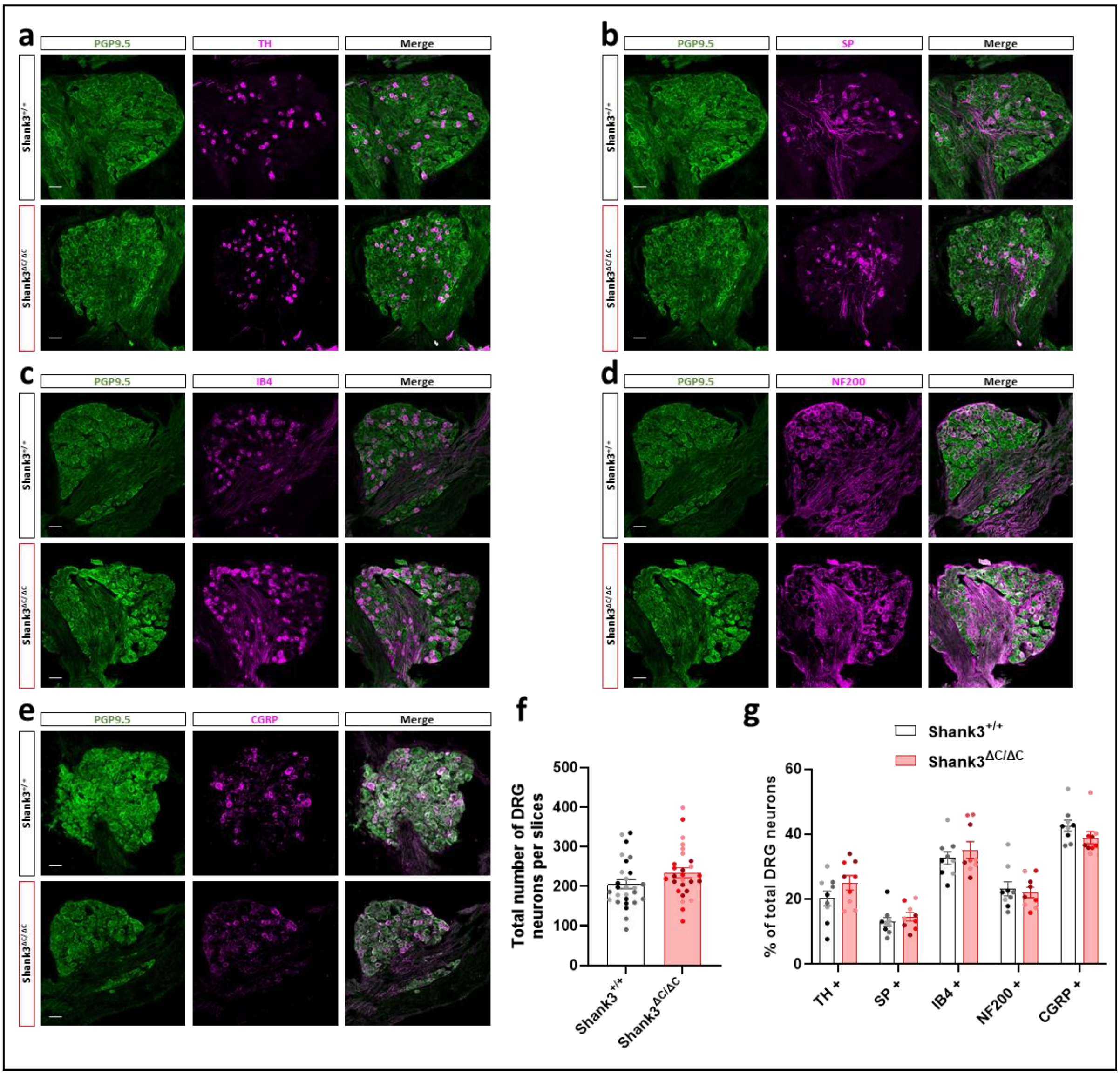

**Supplementary Fig. 5**

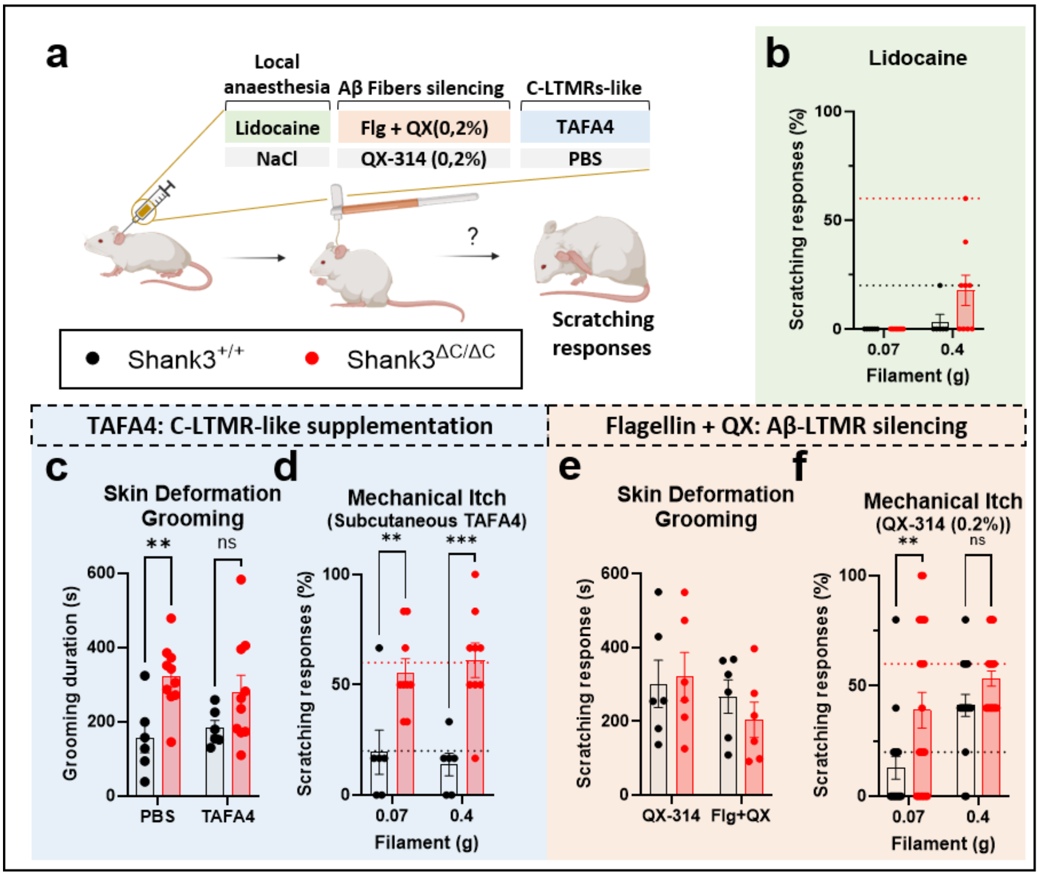

**Supplementary Fig. 6**

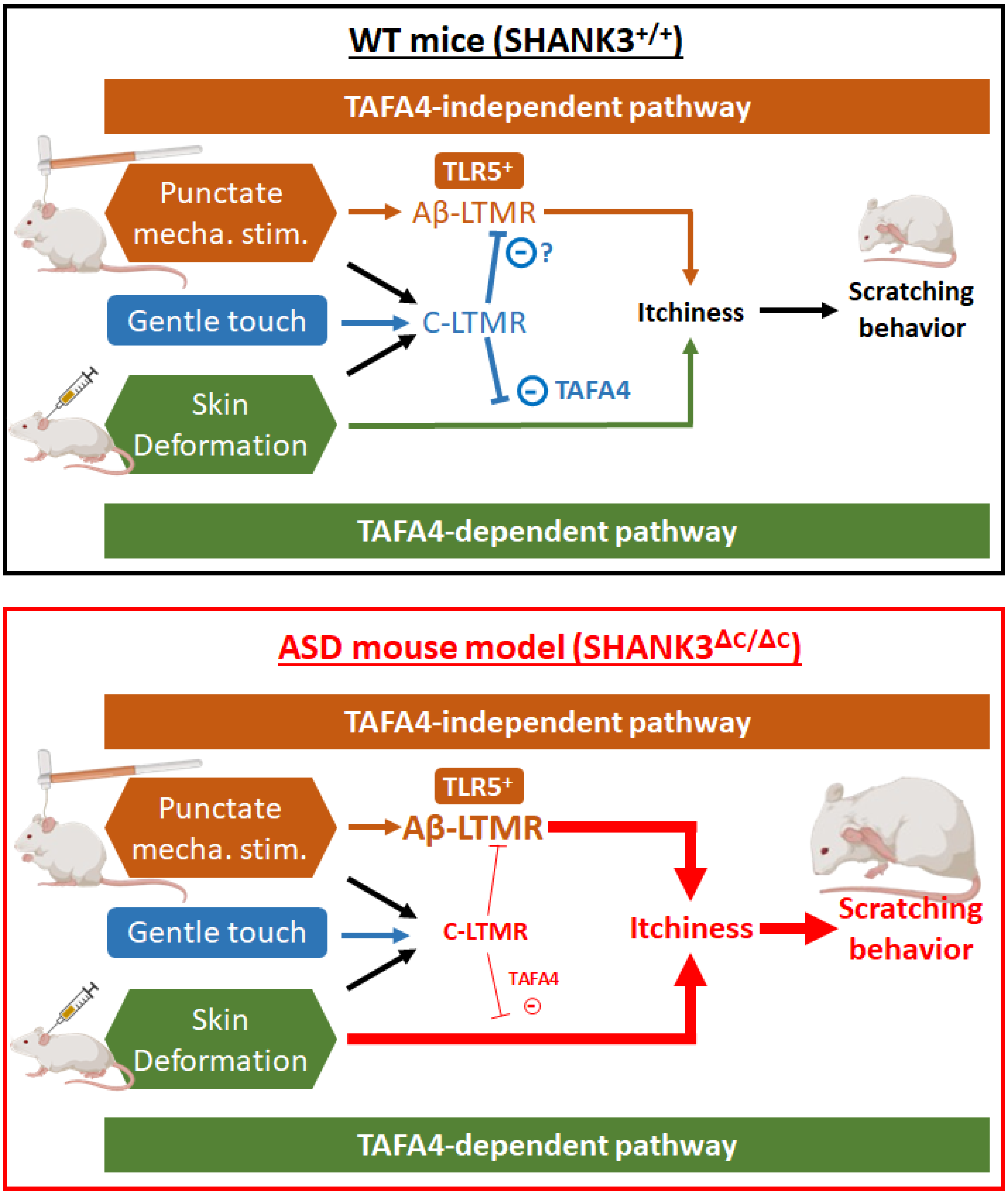
